# Display of clustered antigen by follicular dendritic cells tunes B-cell receptor activation

**DOI:** 10.1101/2025.11.05.686827

**Authors:** Ali Shahrokhtash, Hiroaki Ogasawara, Kristian Savstrup Kastberg, Khalid Salaita, Søren E. Degn, Duncan S. Sutherland

**Author notes:** Corresponding authors Correspondence to Søren E. Degn or Duncan S. Sutherland.

## Abstract

To generate high-affinity antibodies against pathogens, B cells engage antigens displayed on the surface of follicular dendritic cells (FDCs) in secondary lymphoid tissues. Binding of cognate antigen by the B-cell receptor (BCR) causes Src kinase activation and phosphorylation of SYK (pSYK), serving as a platform for downstream signaling. However, how presentation of antigen by FDCs influence antigen binding and intracellular signaling in B cells remains unclear. Here, we show that antigen occurs in submicron pre-clusters on FDCs. We use nanoscale ligand patterning to present clustered antigens on tension force tethers in relevant cluster geometries in vitro. Clustering of antigens greatly increases BCR activation in a pattern-size-dependent manner, correlating to exclusion of the tyrosine phosphatase CD45 from BCR-antigen complexes. Increased height of antigen placement reduces CD45 exclusion with concomitant reduction in pSYK. Super-resolution imaging reveals reduced levels of pSYK at the edges of antigen patterns where CD45 interactions are greater, indicating a geometric control of BCR triggering.

## Main

Antibody production by B cells is a fundamental component of the adaptive immune response^1^. Triggering of the B-cell receptor (BCR) by a cognate antigen leads to BCR signaling, ultimately resulting in the production of class-switched (IgG, IgA, and IgE) and affinity-matured antibodies^1^. Immediately following BCR triggering, Src family tyrosine kinases (Lyn, Fyn, and Blk) phosphorylate immunoreceptor tyrosine-based activation motif (ITAM) domains of the BCR complex, leading to the recruitment and phosphorylation of the spleen tyrosine kinase (SYK)^2^. Phosphorylated SYK (pSYK) in turn recruits and activates further adaptor proteins (e.g. BLNK and PLCγ) propagating BCR signaling. BCR signaling is controlled by membrane-bound (e.g., CD45^3^) and cytosolic phosphatases (e.g., SHP and SHIP^4^). Despite recent structural insights into the BCR complex^5–7^ and a detailed understanding of the signaling cascade, the mechanism by which antigen binding triggers a BCR signal remains unclear^8^. Over the last decades multiple models have been proposed to explain how antigen binding could translate to BCR signaling through changes in ITAM localization (cross-linking or clustering^9^, conformation-induced oligomerization^10^), ITAM accessibility (dissociation-activation^11^), or kinase/phosphatase balance (kinetic segregation^12^); however, the molecular mechanism remains unresolved.

BCR activation can accommodate antigens of very different sizes and valencies, from individual proteins or protein fragments to viral particles or bacteria^8^. However, B cells also commonly encounter antigens retained in intact forms within immune complexes (ICs) on the surface of follicular dendritic cells (FDCs)^13^. Current biophysical models used to study antigen engagement and BCR activation at membrane-membrane contacts are based on antigens presented at supported lipid bilayers^14^ and do not address the size of the antigen-presenting complexes. A number of recent studies have shown the potential for nanoscale ligand patterning to study cellular adhesion and signaling in vitro^15,16^.

Here, we first determine in vivo antigen cluster sizes on FDCs and then develop a robust in vitro nanopatterning platform to present antigens unclustered or preclustered in systemically varied size domains across the physiological range. We integrate DNA tension probes into the antigen presentation to quantify mechanical events upon BCR engagement. We find that B cells respond more efficiently to nanoclustered antigens as opposed to non-patterned antigen surfaces. This effect is pattern-size-dependent and is driven by CD45 exclusion. Here, CD45 exclusion and concomitant BCR activation were reduced as the antigen height was increased. We observe that the size of presented antigen clusters from FDCs modulated the B-cell response with a 200-300 nm antigen cluster size window providing optimal physiological efficiency. In summary, we uncover a new operating model for how B cells sense antigens via their BCRs, based on antigen-driven exclusion of bulky extracellular phosphatases.

### Antigens are presented as nanoscale clusters on FDCs

Although B cells can respond to soluble monovalent antigens, and to a greater extent, soluble multivalent antigens^12,17^, fewer and less-affine interactions are required for B-cell activation by membrane-scaffolded antigens^18^. In a physiological context, B cells commonly encounter antigens displayed by FDCs in follicles^13^. The follicle presents an environment with low proteolytic activity^19^, FDCs serve as dynamic antigen libraries by retaining antigen in recycling endosomes^20,21^, and the spatial organization of the FDC network serves to further control the long-term retention of antigen in GCs^8^. Taken together, this suggests that a highly orchestrated presentation of intact antigen is critical to efficient GC responses. Indeed, early ex vivo observations indicated that antigens are presented on dendrites in a highly ordered pattern^19^, suggesting a periodic arrangement of binding sites^22^. In vivo, these immune-complex coated bodies (iccosomes) were found to measure 300-700 nm in diameter^23^. However, a functional relevance of this observation was never established.

To investigate this phenomenon, we immunized mice with ovalbumin (Ova) in alum intraperitoneally, and intravenously administered fluorescently labeled Ova 7-13 weeks later to load splenic FDCs with fluorescent immune complexes (supplementary Fig. S1). The next day, the spleen was harvested, sections were prepared and counterstained for CD169 (MOMA-1) to define the marginal zone, IgD to visualize naïve B cells, and CD35 (CR1) to identify FDCs, then analyzed via Airyscan super-resolution microscopy. Within the IgD^+^ zone of the follicle, which is bounded by CD169^+^ marginal zone macrophages, fluorescently labeled Ova was observed to decorate the reticular FDC network in discrete antigen clusters (Fig. 1a). The quantification of nearly 3,000 clusters revealed a size distribution from ∼100-600 nm, peaking at ∼150-350 nm (Fig. 1b).

**Figure 1:**
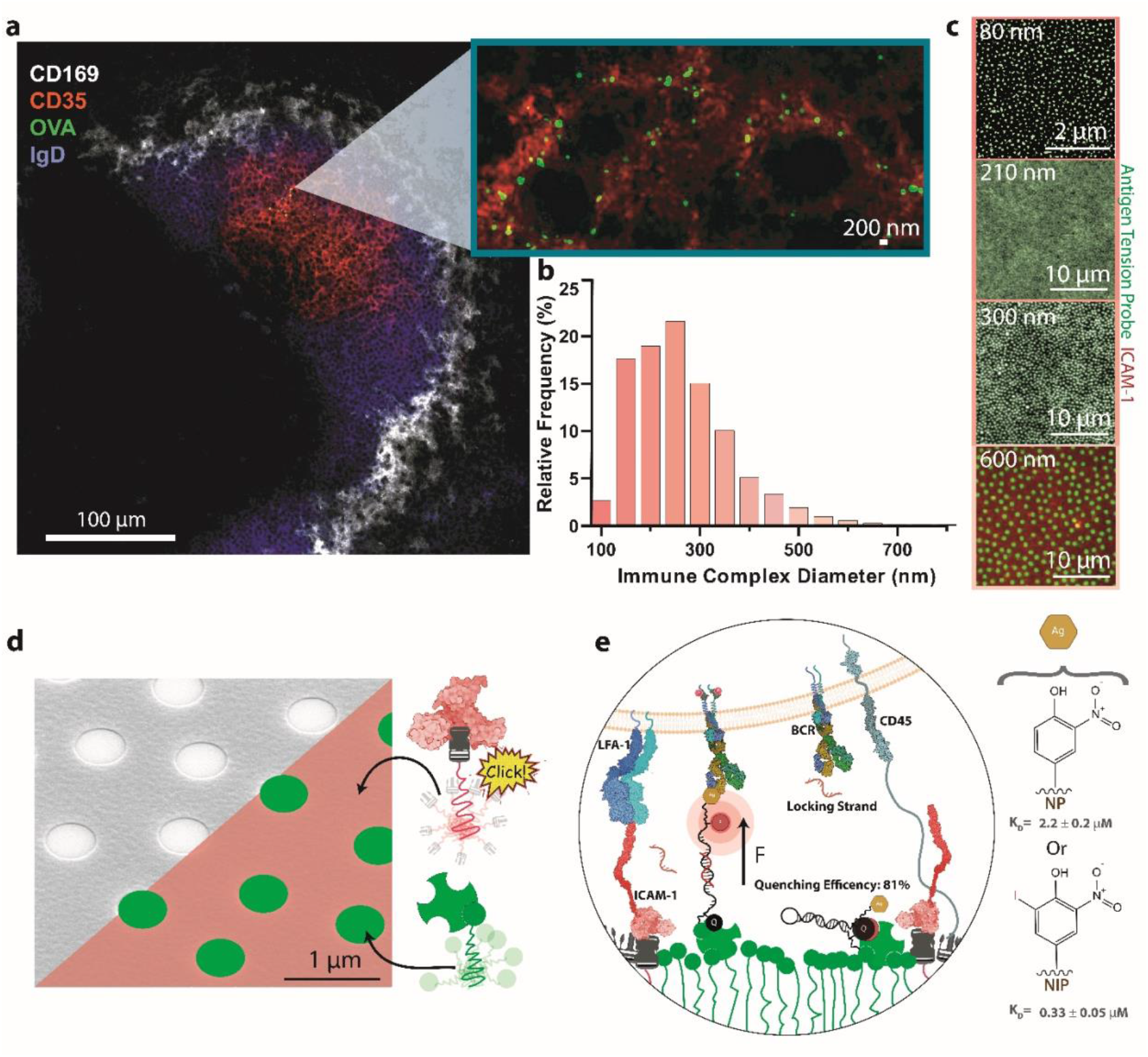
Antigens form nanoscale clusters in vivo which can be mimicked via nanofabrication of biomolecular assemblies. **a**, Representative fluorescence image of a splenic follicle containing CD35^+^ follicular dendritic cells (FDCs, red) presenting ovalbumin antigen (green), and naïve B cells (blue). The inset shows a magnified view, revealing antigen clusters on the FDC network. **b**, Quantification of antigen cluster sizes. The histogram shows the frequency distribution of antigen cluster diameters (n=4 animals, n=2,895 clusters). The bin width is 50 nm. **c**, Representative images of nanofabricated clusters, measuring 80 nm (super-resolution DNA-PAINT), 210 nm (TIRFM), 300 nm (TIRFM), and 600 nm (CLSM). **d**, Tilt-view scanning electron microscopy (SEM) micrograph with a partial pseudocolor overlay indicating the patterning of anti-fouling biospecific PEG brushes. The substrate was generated via sparse colloidal lithography and deposition of a thin sacrificial chromium (Cr) layer. Incubation with the first polymer brush (green), followed by wet etching of the sacrificial Cr layer, and subsequent incubation with the second biospecific polymer brush (red) yielded a fully PEGylated and metal-free surface. The polymer brushes are terminated with biotin for binding to avidin-family proteins and azide groups for copper-free click chemistry. Streptavidin is depicted in green and Protein A in red. The Protein A crystal structure was obtained from PDB entry 4WWI **e**, Side-view schematic zoom-in of a naïve B cell interacting with the nanopatterned surface. Upon interaction of the BCR with the antigen conjugated to DNA tension probes, the stem-loop hairpin is mechanically melted by the pulling forces generated by the BCR. This separation of the quencher and fluorophore results in an increase in fluorescence intensity, reporting a mechanical event above the ∼6 pN threshold. When the probe is under mechanical force, a locking strand can anneal to it and keep it open, allowing for the accumulation of mechanical events. The model antigen system used includes the haptens 4-hydroxy-3-nitrophenyl (NP) and 4-hydroxy-3-iodo-5-nitrophenyl (NIP). Protein structures were obtained from PDB: IgM BCR (7XQ8), LFA-1 (5ES4), CD45 (1YGR and 5FMV), FcγRIIb (3WJJ), IgG1 (1IGY), ovalbumin (1OVA), CD19/CD81 (7JIC), and CD21 (2GSX). The ICAM-1 structure was generated via AlphaFold 3.

Having confirmed a relevant size scale for FDC-displayed antigen at the nanoscale, we next asked what the functional impact is for B-cell activation. To this end, we applied nanofabrication to mimic in vivo antigen cluster sizes presentation in vitro (Fig. 1c)^24^. We leveraged a novel biomolecule nanopatterning platform that generates nanopatterns of two biomolecules on a transparent, passivated glass substrate^25^. Here sparse colloidal lithography^24^ (SCL)-prepared thin sacrificial Cr layers directed covalent attachment of passivating antifouling polymer brushes. We assembled PAcrAm-g-PEG-biotin into circular nanoscale domains and coated the background with PAcrAm-g-PEG-N_3_ after Cr layer etching to produce a fully PEGylated surface with excellent antifouling properties and two nanopatterned orthogonal biospecific tags (Fig. 1d)^25^. We used biotin tags to immobilize Streptavidin (SA) as a linker for biotinylated proteins/DNA and azide tags (N_3_) to enable copper-free strain-promoted alkyne-azide cycloaddition (SPAAC) click chemistry to covalently immobilize DBCO-tagged biomolecules (Fig. 1c and 1d).

B cells exert mechanical forces during affinity discrimination and antigen uptake^14,26–29^, therefore, we integrated DNA-based molecular tension probes that respond rapidly to external forces^30^ conjugated to small (<1 kDa) hapten antigens as force threshold sensors and patterned these into clusters of 80 to 600 nm in diameter via biotin-SA interactions. We ensured a constant global antigen coverage of ∼3,500 antigens/μm^2^ via quantitative fluorescence microscopy^31^ with ∼20% nanopattern coverage (Supplementary Figs. S2-3).

The tension probes were designed as 25-mer phosphorothioate-modified, nuclease-resistant DNA (Supplementary Fig. S4-6) forming a stem-loop hairpin^32^. The force threshold for a 50% probability of melting (F_1/2_) measured ∼ 6 pN (Supplementary Information). The probes contained a Cy3B fluorophore which was efficiently quenched by a BHQ2 quencher in the closed stem-loop conformation. The opening of the stem-loop under mechanical force during BCR engagement separates the fluorophore and quencher serving as a light-up reporter of mechanical events (Fig. 1e) above the force threshold (F_1/2_ ∼ 6 pN) (Supplementary Fig. S5 & Supplementary Information). The hairpin loop recloses when the force drops below F_1/2_ allowing observation of dynamic mechanical events on the surface (Supplementary Fig. S7 & Supplementary Video S1). To capture rapid transient events we used a locking strategy^33^ based on the inclusion of a 17-mer DNA locking strand that partially complements a cryptic binding site exposed under mechanical force to keep the stem-loop open after force cessation (Supplementary Fig. S8).

In the background, we provided ICAM-1-Fc bound to DBCO-tagged Protein A to mimic ICAM-1 on the surface of FDCs^34^ which enhances cellular adhesion and lowers the activation threshold of B cells via LFA-1^35^. Control non-patterned surfaces were prepared where antigen and ICAM-1-Fc are co-presented homogeneously with matched global antigen density.

To systematically study FDC–B cell interactions, we employed the B1-8i heavy chain knock-in J-kappa knock-out model^36^, in which approximately 60% of naïve B cells carry a BCR with affinity for the haptens 4-hydroxy-3-nitrophenylacetyl (NP) and 4-hydroxy-3-iodo-5-nitrophenylacetyl (NIP)^12^ in the low micromolar range (NP: ∼2.2 ± 0.2 μM; NIP: 0.33 ± 0.05 μM)^26^.

### B-cell activation depends on the nanoscale antigen cluster size

We monitored cell spreading and early BCR activation at nano-clustered antigen patterns (80 nm, 200 nm, 300 nm, and 600 nm) and non-patterned homogeneous surface culture wells after 10 minutes. We observed mechanical events from the cell areas across all cluster sizes (Fig. 2a & Supplementary Fig. S9). For pattern sizes above the diffraction limit, the signals clearly originated from the antigen clusters.

**Figure 2:**
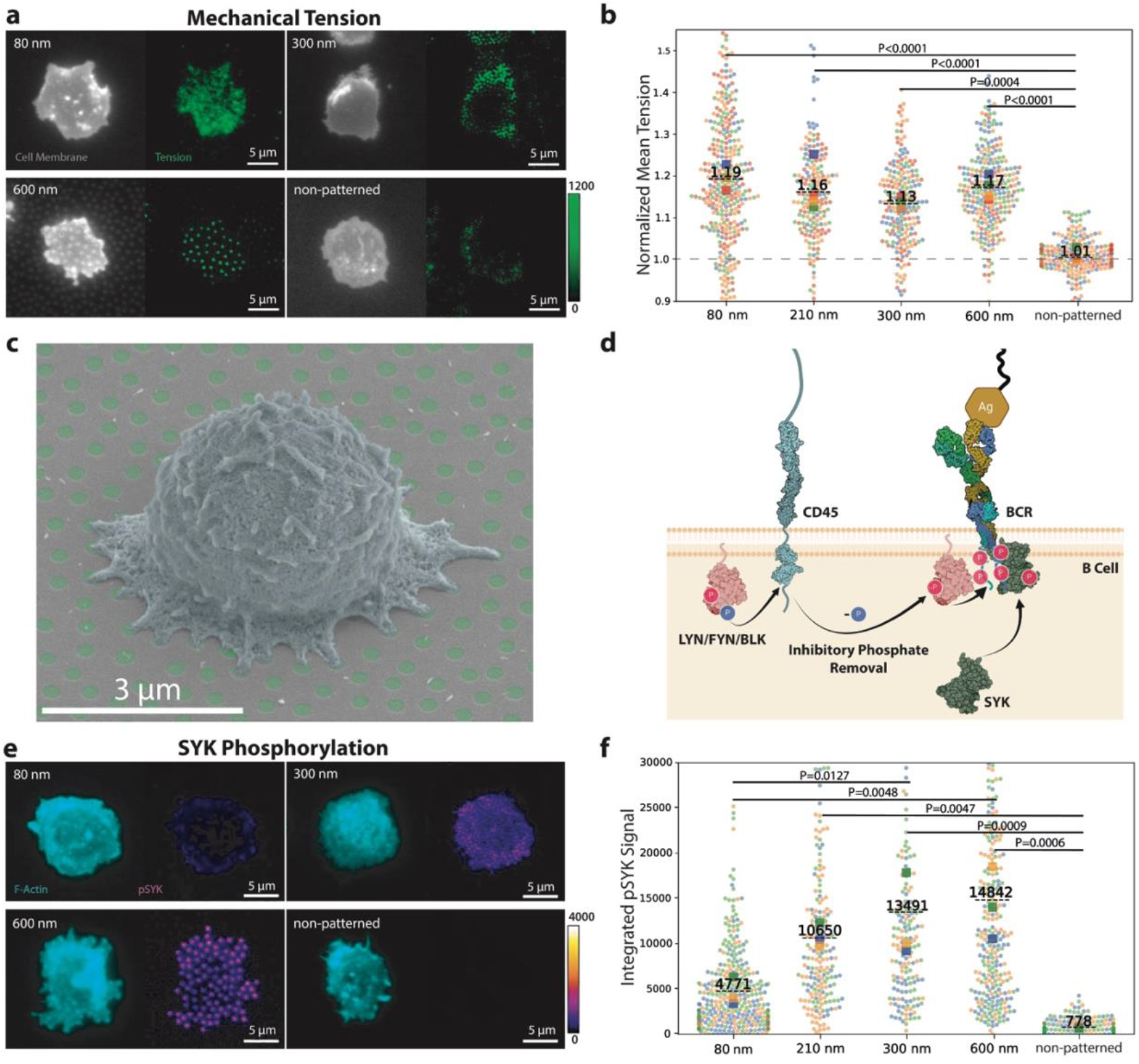
Naïve B-cell activation on NP antigen nanopatterns. **a**, Representative TIRFM images of the cell membrane (gray) and locked mechanical tension signal (green), captured 10 minutes after seeding on the surface. **b**, Quantification of mechanical events across all nanopattern sizes. All surfaces were prepared to have the same global antigen density. Each circle represents the background-subtracted mean signal of a cell, color-coded by experimental repeat (n = 4 animals); squares represent the mean of each biological replicate, and the dashed line represents the overall mean across all biological replicates. **c**, SEM image of a naïve B cell interacting with 300 nm NIP antigen cluster patterns. The micrograph is false-colored to enhance visibility, with the patterns in green and the cell in blue. **d**, Schematic representation of the initial BCR activation steps. The phosphatase CD45 is required to dephosphorylate the inhibitory phosphate on Src family kinases such as Lyn to prime them for activation. Activated Lyn kinase phosphorylates the immunoreceptor tyrosine-based activation motifs (ITAMs) of the BCR complex, allowing the docking of the kinase SYK to the ITAMs, which results in autophosphorylation of SYK. At this stage, downstream B-cell signaling is initiated. **e**, Representative pseudocolored images of F-actin (cyan) and phosphorylated SYK (pSYK) signal (purple). **f**, Quantification of the background-subtracted integrated pSYK signal (n = 3 animals). *P*-values were determined by one-way ANOVA with Tukey’s multiple comparison test between experimental replicate means

The cell spread areas were comparable for the low-affinity NP antigen (∼ 50 μm^2^) and NIP antigen (∼ 60 μm^2^) (Supplementary Figs. S10-11). The mean tension (mechanical events above ∼ 6 pN), normalized to the background, was significantly higher on nanopatterned substrates than on non-patterned surfaces (Fig. 2b) and increased with the high-affinity NIP antigen (Supplementary Figs. S12-13). Notably, few mechanical events occurred on the non-patterned surfaces (NP 1.01 < NIP 1.11). Taken together, these findings indicate that antigen affinity as well as clustering play a role in the mechanical response of the BCR.

B cells showed extended filopodia to probe antigen patterns under both TIRFM and high-resolution SEM imaging (Fig. 2c and e, and Supplementary Fig. S14). This agreed well with recent 4D imaging using lattice light-sheet microscopy which suggested that dynamic topographical features of the cell surface facilitate antigen screening by B cells^37^.

Having confirmed specific interactions with nanopatterned antigens and similar mechanical engagement across pattern sizes, we investigated the early activation events in naïve B cells. Antigen engagement in B cells triggers intracellular signaling cascades mediated by Src family kinases (such as Lyn) and CD45^38^ leading to the phosphorylation of ITAMs on BCR-associated Igα and Igβ chains^39^ which activate SYK, resulting in the formation of pSYK clusters that act as signaling hubs for downstream BCR signaling^40^ (Fig. 2d).

Accordingly, we quantified pSYK as a readout of BCR activation. The pSYK signals co-localized with the antigen patterns (see Fig. 2e). We measured integrated pSYK signals per cell after background subtraction (Fig. 2f). Non-patterned surfaces showed the lowest pSYK signals, followed by the 80 nm patterns, with a plateau at 300 nm clusters. The same trend was observed for the NIP antigen (Supplementary Figs. S15-16). Altering the antigen density from 20% (to 12– 27%) did not affect this trend and the pattern-size-dependent activation threshold persisted (Supplementary Fig. S17).

Comparing these results with the in vivo dimensions and relative abundance of antigen clusters on FDCs (Fig. 1c) suggested that antigen clusters of 200–300 nm in diameter are the most physiologically efficient for B-cell activation.

### CD45 exclusion drives B-cell activation at the nanoscale

We explored the potential role of CD45 in regulating the observed antigen cluster size dependency of BCR triggering.

CD45 is abundantly expressed on B cells, with the long form dominating on naïve mature B cells. CD45 plays a dual role in B-cell activation, enabling the initiation of signaling cascades indirectly through dephosphorylation of the inhibitory tyrosine residues of Src family kinases^38,41^, but also directly attenuating activation by dephosphorylating activating phosphotyrosines on Src kinases, ITAMs, and SYK, thus maintaining homeostasis^42^. In T cells, the kinetic segregation model^43^ posits that activation is regulated by spatial segregation on the basis of molecular size within the immunological synapse. Larger molecules like CD45 are excluded from close-contact regions formed by TCR–peptide-MHC interactions, creating microenvironments where smaller activating molecules can phosphorylate signaling motifs. The absence of CD45 in these regions reduces its inhibitory phosphatase effect, allowing stable propagation of activation signals.

Here, we observed close contacts between B-cell membranes and antigen patterns (indicated by high Reflectance Contrast Interference Microscopy (RICM) signals Supplementary Fig. S17b). We investigated CD45 localization in relation to these contacts observing clear exclusion of CD45 from the regions occupied by antigen nanopatterns (for the 300 nm and 600 nm patterns which exceed the diffraction limit). Figure 3a & Supplementary Fig. S18 show CD45 immunostaining via TIRFM after 10 minutes of naïve B-cell interaction with NP/NIP antigen clusters.

**Figure 3:**
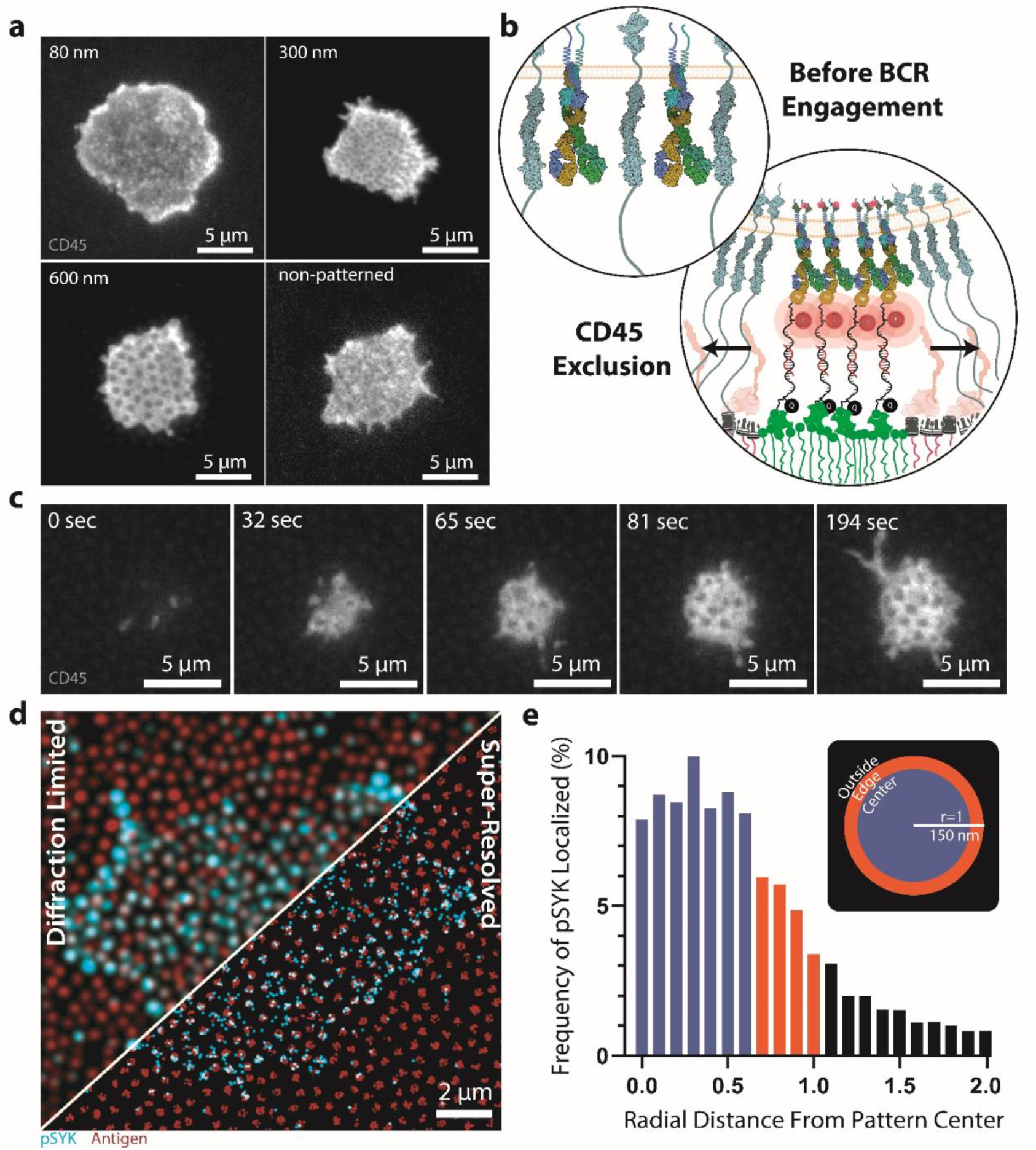
CD45 is excluded upon BCR engagement. **a**, Representative TIRFM micrographs of CD45 (gray) staining in B cells 10 minutes after engagement with the NP nanopatterns. **b**, Schematic illustration of CD45 exclusion upon BCR engagement. Before BCR engagement, BCRs and CD45 are intermixed; however, upon engagement with antigens, BCRs cluster together, exerting mechanical pulling forces, and CD45 is rapidly excluded. **c**, Time-trace of live-cell CD45 exclusion upon engagement with 600 nm antigen patterns (see also supplementary movies 2-3). **d**, Diffraction-limited and DNA-PAINT super-resolved images of phosphorylated SYK (pSYK, cyan) and 300 nm antigen nanopatterns (red). **e**, Probability of localizing pSYK as a function of distance from the center of a nanopattern, measured based on super-resolution DNA-PAINT imaging of NIP antigen patterns and pSYK clusters. 75% of pSYK localizations were identified to be at least 20 nm away from the edge of the nanopatterned antigen clusters. The blue bars indicate the central region of the patterns, orange the edge region, and black bars indicate the pSYK molecules localized outside of the antigen-cluster nanopatterns. (n = 5 cells, 3,436 pSYK clusters total).

CD45 was excluded from BCR clusters but remained between them to regulate homeostasis (Fig. 3b). This aligns with the kinetic segregation model for T cells and observations by Shelby et al., where the selective exclusion of membrane proteins like CD45 from BCR clusters, based on differential partitioning into liquid-ordered nanodomains, drives signaling^44^.

To understand the dynamics of this process, we recorded the exclusion dynamics of CD45 using fluorescently labeled primary monoclonal CD45 antibodies for live-cell imaging on 600 nm nanopatterns, which are easily resolvable via TIRFM. Within 30 seconds of surface engagement, CD45 exclusion from antigen nanopatterns was evident (Fig. 3c and Supplementary Movies 2–3).

The exclusion of CD45 and resultant reduction in dephosphorylation of pSYK/ITAMs near nanopatterned antigen clusters may underlie the observed pattern-size-dependent BCR triggering (Fig. 2f). Smaller patterns have larger relative perimeters where CD45 and pSYK may interact ( ∼4 times larger circumference/area for 80 nm compared to 300 nm patterns), implying that larger patterns may better shield downstream signaling components allowing more robust signal propagation.

To further investigate the interactions at the antigen nanopattern periphery, we employed DNA-PAINT super-resolution microscopy^45,46^ on 300 nm antigen nanopatterns (Fig. 3d). We localized more than 3000 pSYK clusters and compared their positions relative to the antigen nanopatterns and plotted their radial distance distribution (Fig. 3e). Importantly, a sharply reduced level of pSYK was observed at the edge of the patterns (within 50 nm of the edge) than at the center (Fig. 3e).

These findings, combined with the pattern-size-dependent activation of naïve B cells, indicate the likely involvement of CD45. Here CD45 access to dephosphorylate pSYK/ITAMs at BCR sites is reduced by exclusion from antigen patterns by membrane proximity, leading to increased activation at patterned surfaces compared to non-patterned. We observed a reduction of pSYK at the outer edge of the patterns, likely from partial access of CD45 to pSYK, by the molecular reach or diffusion of pSYK/BCR complexes. In the case of 80 nm clusters, this likely affects the whole patterned region, whereas, for larger patterns, it affects only a small fraction of the pSYK/BCR complexes, resulting in higher activation at larger patterns, as shown in Fig. 2f.

### Matching antigen height to CD45 reduces B-cell activation

In the T-cell kinetic segregation model, the size of the CD45 ectodomain, up to 50 nm^47^, is suggested to critically impact its exclusion from the signalosome.

To better understand how CD45 exclusion affects BCR activation by nanoscale antigen clusters, we kept the antigen density constant but varied the height of the antigen probes from 5 nm to 48 nm, comparable to the dimensions of the CD45 ectodomain, with the hypothesis that a taller antigen would result in less CD45 exclusion, thus modulating activation. This was achieved by varying the length of DNA probes of identical chemistry, carefully designed to avoid undesired secondary structures, presented on 300 nm antigen nanopatterns.

The antigen conjugated to the standard DNA tension probe (approximately 18 nm in height when unfolded, including linkers and SA, Fig. 4a), gave a high level of pSYK signal (Fig. 4b). In contrast, measurements with extended antigens up to 48 nm (Fig. 4c) yielded substantially lower pSYK signals (Fig. 4d). We compared pSYK levels across 5 different antigen probe heights, showing a significant reduction in BCR triggering with increasing height (Fig. 4e) and low but non-zero pSYK levels for the tallest antigens.

**Figure 4:**
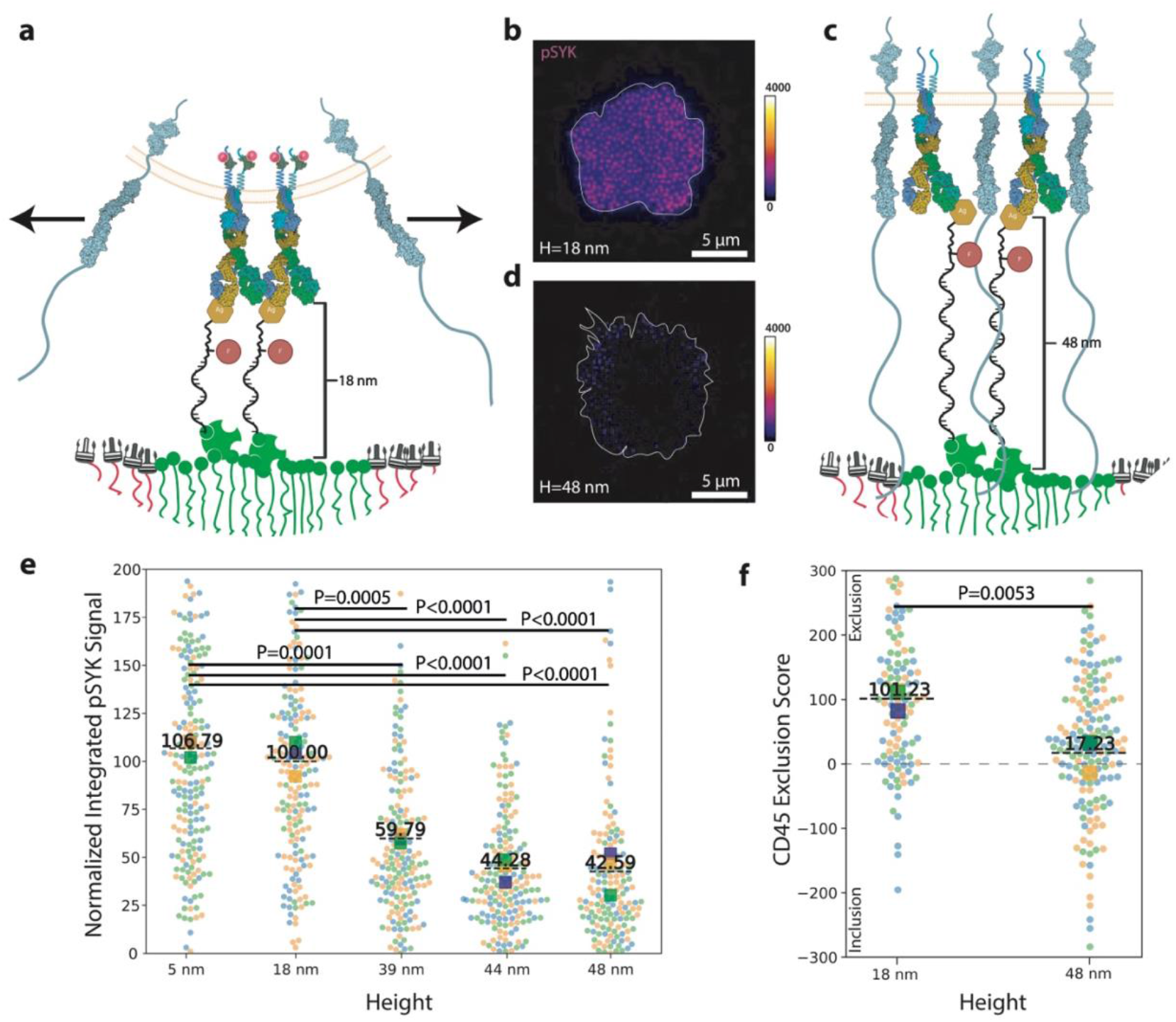
Antigen height modulates CD45 exclusion and B-cell activation. **a**, Schematic illustrating that BCR engagement with the normal short antigen (which extends under mechanical force to approximately 18 nm) should result in the exclusion of CD45 due to a size mismatch. **b**, Representative image of the pSYK signal in a naïve B cell after 10 minutes of interaction with the 18 nm-tall NP-conjugated antigens on 300 nm patterns. The white line indicates the cell border. **c**, Schematic illustrating that when the antigen height (48 nm) is comparable to the size of the extracellular domain of CD45, CD45 exclusion should be less effective. **d**, Representative image of the pSYK signal in a naïve B cell after 10 minutes of interaction with the 48 nm-tall NP-conjugated antigens on 300 nm patterns. The white line indicates the cell border. **e**, pSYK signal at various antigen heights on 300 nm NP patterns. **f**, CD45 exclusion scores for the normal 22% GC NP probe (18 nm extended) and the long 76-mer ssDNA probe (48 nm) on 300 nm patterns. Each circle represents a normalized CD45 exclusion score for an individual cell, calculated by comparing the mean CD45 signal in regions with underlying antigen nanopatterns to the remaining spread area of the cell. Positive values indicate the exclusion of CD45 from the antigen nanopatterns (on average), while negative values indicate a lack of CD45 exclusion. The circles in panels **e** and **f** are color-coded by experimental repeats (n = 3 animals); the squares represent the mean of each biological replicate, and the dashed line represents the overall mean across all biological replicates. P-values were determined by t-test (for **f**) and one-way ANOVA with Tukey’s multiple comparison test (for **e**) on the experimental replicate means.

We quantified the level of exclusion at 18 nm or 48 nm tall antigens at 300 nm patterns by comparing the CD45 signals within patterns to the mean intensity outside the patterns but underneath the cells, with a negative score indicating no exclusion (Fig. 4f). The high levels of pSYK with the 18 nm probe correlated with high CD45 exclusion, whereas the extended antigen showed both lower pSYK and lower CD45 exclusion. We used the force probes to compare antigen engagement for short (18 nm) and long (48 nm) antigens. Here, the number of mechanical events was not significantly different for the ∼ 6 pN force probes, indicating that there was similar accessibility of the antigen and similar BCR engagement (Supplementary Fig. S19).

Taken together, these findings indicate that CD45 exclusion directly correlates with pSYK activity for clusters with the same antigen density, cluster size, and level of BCR mechanical engagement.

## Conclusions

In this work, we found that antigen immune complexes are presented as sub-micron clusters of 150-350 nm in size by FDCs in germinal centers. Nanoscale patterning of antigen in vitro was used to mimic the FDC surface-presentation and explore the role of antigen complex size on BCR activation as well as forces exerted on the antigen by the B cells. We confirmed that B cells spread on these surfaces, interact with the patterns, and exert tugging forces on the antigens. Importantly, the mean normalized tension was comparable between antigen nanopatterns from 80 nm to 600 nm, and significantly higher than on non-patterned surfaces. The amount of pSYK, a key component of the BCR-proximal signaling hub driving downstream signal transduction, was comparable on the 210 nm to 600 nm patterns, but significantly lower on the 80 nm and non-patterned surfaces. Taken together, this indicates a sweet spot for the BCR-activating capacity of antigen cluster dimensions matching the range of antigen immune complex sizes observed on FDCs in vivo. Prior insights into the physiology of FDC antigen presentation have highlighted 1) the importance of the low proteolytic activity in the follicle in maintaining antigens in intact form^19^, 2) the capacity of FDCs to internalize immune complexes in recycling endosomes^21^, and 3) the ability of the FDC network to create a gradient of antigen availability^48^, as key factors in the longevity of antigen deposits to support long-lived germinal centers for efficient affinity maturation. Our findings suggest that the organization of immune complexes into defined antigen patches is a fourth central contributor to the extraordinary capacity of FDCs to support efficient B-cell responses. Indeed, highly orchestrated antigen arrays were observed early on by others^49,50^, but whether these highly organized antigen patches are a byproduct of the endosomal recycling mechanism or are orchestrated by other processes and cytoskeletal forces remains to be determined. No matter their origin, the functional implications of these antigen clusters was never determined.

Here, we investigated the importance of antigen clustering for BCR activation, shedding new light on the mechanism of BCR triggering^8^. Overall, our findings provide direct evidence of a link between CD45 exclusion and the transduction of antigen binding events into activated BCR complexes, which supports the extension of the T-cell kinetic segregation model to BCR triggering, which has been termed the antigen footprint model^8,12^. In that model, based on studies with soluble monovalent and multivalent model antigens, it was proposed that the footprint of the bound antigen causes a subtle and dynamic exclusion of CD45 across the global receptor population engaged by antigen. However, in that study, although baseline BCR localization was defined using DNA-PAINT super-resolution microscopy, the potential dynamic redistribution of BCRs and the localization of CD45 were not addressed. Here, we present direct evidence that BCR engagement, CD45 exclusion, and B-cell activation are intimately linked. The degree of SYK phosphorylation is correlated with the distance from the perimeter of nanopatterns and anti-correlated with the proximity to CD45, effects that can be shortcircuited when the antigen presentation height is extended sufficiently to accommodate CD45 within the patterns.

In this perspective, it is of interest to consider the relative dimensions of the key receptors involved in antigen retention on the FDC surface, FcγRIIb–IgG molecule complexes and complement receptor-complement fragment complexes. The former are quite compact, whereas at least the complement receptors themselves are rather extended, but may wrap around physiological target protein-complexes. Moreover, for larger immune complexes or intact particulate antigens such as bacteria or viruses, the antigenic surfaces themselves might be envisioned to form the appropriate nanopattern platforms for activation. Indeed, the observed sweet spot for pattern size is on the order of many relevant viral particles (influenza ∼100 nm, HIV 100-120 nm, measles 100-300 nm, RSV 150-300 nm), which typically present highly repeating and efficient arrays of surface antigens that are emulated by VLP and DNA-origami-based vaccine platforms^51–54^. The structured presentation of antigens in immune complex clusters may allow FDCs to similarly mimic such naturally strong activators of humoral immunity. Notably, the exclusion should be dramatically more efficient because it is potentiated by membrane-membrane (or membrane-surface) approximation. The effect of antigen binding on the kinetic segregation of BCR and phosphatase activities would also be envisioned to be directly dependent on the total docking time, given that fast dissociation of the antigen would allow the BCR to quickly meet phosphatases and thus revert to a resting state. Again, this consideration would favor BCR activation by membrane-scaffolded antigen. Local depletion of signal components downstream of the BCR could, however, be imagined to set a limit for individual receptor activity. Moreover the long-term paucity of CD45 phosphatase activity likely causes signal attenuation because the inhibitory phosphatase activity of Csk would no longer be mitigated. Altogether, this establishes a framework for the periodicity of antigen-driven BCR signaling entailed in the prevailing model for B-cell activation and GC B-cell recycling, that may inform future vaccine designs.

Importantly, there are potential caveats and differences between the present system and commonly employed supported lipid bilayer (SLB) approaches. Firstly, inherent to their ability to present stable nanopatterns, our surfaces display no lateral mobility. However, this may actually better represent the FDC scaffolded antigen than freely diffusible SLBs, both because stable immune complexes are likely largely static and because FDC-displayed antigens may, in fact, be anchored by the cytoskeleton, as suggested by the highly ordered periodic arrangement observed by others^55^. Secondly, the tension signal quantified by the antigen force gauge tether approach represents the accumulated ‘history’ of mechanical events of the cell. As noted previously for similar TCR– pMHC measurements, this means that the tension signal depends on the mechanical activity of the B-cell cytoskeleton, engagement with BCRs, and on BCR copy number^56^. However, in the present study these factors were either controlled (BCR copy number, BCR engagement affinity) or an inherent quality of the readout (tension signal and cytoskeletal activity), hence mitigating such concerns. Third, our setup did not enable early temporal resolution sufficient to permit the assessment of ultrafast events occurring upon initial antigen contact. This is difficult to overcome, firstly because the time between B cell seeding over the surfaces until contact with the antigen patterns is non-uniform, precluding a synchronized starting point for acquisition, secondly because the imaging modalities employed are either limited by time or resolution to sufficiently resolve these phenomena in real-time at the molecular scale.

Here, we specifically investigated antigen and ICAM-1 nanopatterning for the reasons stated at the outset, but given the versatility of our approach, the strategy has broad applications in studying signaling in immunity and other cellular settings. For B cell–FDC interactions, this could include studies of patterning auxiliary integrin adhesion molecule interactions (VLA-4–VCAM-1), B-cell co-receptor complex interactions with complement (CD19/CD21/CD81–C3d), and B-cell inhibitory Siglec interactions with glycosylated cell surface molecules (CD23–glycans). Extending this last point, immunoreceptor tyrosine-based inhibitory motif (ITIM)-containing receptors strongly suppress the signal propagation from the BCR that is triggered by low-valency antigens, and physical exclusion of these receptors from the BCR may be equally important to CD45 exclusion for the efficient coupling of BCR triggering and B-cell activation^8^. Furthermore, we focused on naïve B-cell activation, but other cell states, such as those represented by cycling GC B cells, would be of high relevance to study^51^. Finally, the approach could also be extended to other immune cell subset interactions where spatial receptor patterns may play a key role, such as B–T cell and T–DC interactions.

## Acknowledgments

We would like to thank Joe Mancuso (Emory University) and the Mass Spectrometry Core Facility at the Department of Chemistry, Aarhus University, for assistance with mass spectrometry analysis. We are also grateful to Dr. Alexey Ferapontov, Lisbeth Jensen, Kenneth Green, and Dr. Thomas Wittenborn for their support of animal euthanization. We acknowledge Dr. Layla Pohl for her contributions to the initial two-photon microscopy work and the development of the mice immunization protocol used for in vivo quantification of antigen cluster sizes. This work was supported by the Danish National Research Foundation (DNRF135) and National Institute of Health (NIH, 1RM1GM145394)

## Author information

### Contributions

A.S. designed the study, extracted spleens from animals, purified primary cells, prepared nanopatterned surfaces, conjugated and purified polymers and DNA, conducted all in vitro experiments, analyzed the data, prepared all figures, and wrote the first draft of the manuscript. H.O. contributed to the design of the PS-modified DNA tension probe protocol. K.S.K. performed the in vivo immunization study, including AiryScan super-resolved confocal microscopy. K.S. supervised the DNA tension probe work. S.E.D. and D.S.S. supervised the B-cell nanopatterning experiments and co-authored the manuscript.

## Notes

### Competing Interest Statement

The authors have declared no competing interest.

